# *Arabidopsis thaliana* LSM7 is essential for auxin-mediated regulation of *SAUR* genes and thermomorphogenesis

**DOI:** 10.1101/2023.03.28.534379

**Authors:** Sarah Muniz Nardeli, Vasiliki Zacharaki, Nelson Rojas-Murcia, Silvio Collani, Kai Wang, Martin Bayer, Markus Schmid, Daniela Goretti

## Abstract

Temperature affects plant growth by modulating the expression of genes and subsequent processing of RNAs that govern essential physiological processes. Here, we show that *Arabidopsis thaliana* Sm-like7 (LSM7), a core component of the splicing and decapping machinery, is indispensable for embryogenesis and development. Hypomorphic *lsm7-2* mutants display severe developmental defects that are exacerbated by high temperatures. Transcriptome analysis verified LSM7’s extensive role in gene regulation. In particular, we found that the key regulator of thermomorphogenesis, *PHYTOCHROME INTERACTING FACTOR 4* (*PIF4*), and auxin-related genes, including *SMALL AUXIN UP-REGULATED* (*SAUR*) genes, are misregulated in *lsm7-2*. Auxin metabolic profiling confirmed that auxin homeostasis was disturbed in *lsm7-2*. Importantly, overexpression of the auxin-responsive *SAUR19* gene partially restored thermomorphogenesis defects in *lsm7-2* under high ambient temperature. Taken together, our research provides mechanistic insights into the interplay between RNA processing, hormone homeostasis, and the response to temperature regulation in plants and elucidates LSM7’s essential function in plant temperature acclimation and resilience.

**Significance Statement:** Given their sessile nature, plants cannot escape adverse environmental conditions such as cold or heat. Instead, they continuously adjust their gene expression and RNA processing to regulate growth and physiology in response to their surroundings. In this study, we investigated the role of the core RNA processing factor LSM7 in temperature acclimation in *Arabidopsis thaliana*. We found that LSM7 knockdown mutants were impaired in thermomorphogenesis and, as a result, were hypersensitive to elevated temperatures. At the molecular level, we demonstrated that this temperature sensitivity was caused by the misregulation of key regulators of thermomorphogenesis, including PIF4, auxin homeostasis and signaling, and SAUR genes. Our findings provide valuable insights into the role of RNA processing in plant temperature acclimation.

## Introduction

Temperature is a critical environmental factor affecting plant development. To adjust to environmental changes, plants reprogram their transcriptome by regulating gene expression, RNA processing, and RNA stability. Alternative RNA splicing (AS) is a key mechanism enabling plants to rapidly respond to new environmental conditions(1). Splicing is regulated by the spliceosome, a large ribonucleoprotein machinery, which assembles around heptameric rings of SM and SM-like (LSM) proteins containing U-rich snRNAs. While the SM ring is best known for its function in RNA splicing, LSM rings have a dual function. The nuclear-localized heptameric LSM2-8 ring binds to the 3′ end of the spliceosomal snRNA U6(2) and participates in splicing(3). In contrast, the cytoplasmic LSM1-7 complex participates in the degradation of mRNAs with short poly(A) 3′ ends(4) via the nonsense-mediated decay (NMD) surveillance pathway, thereby protecting the cells from potentially harmful truncated proteins(5). This coupling of AS and NMD enables the cell to co-or post-transcriptionally regulate the level of mRNA(6).

Numerous lines of evidence implicate splicing in plant temperature acclimation(7, 8). However, the contribution of core spliceosome components is still poorly understood. Heat stress has been shown to affect the subcellular localization of several LSM proteins(9) and loss-off LSM5 causes mis-splicing of heat shock transcription factors, resulting in seedling lethality(10). Other studies showed that LSM8 and PORCUPINE (PCP; SME1) are required for acclimation to cold (4°C) and cool ambient temperatures (16°C)(11, 12,13), indicating that (L)SM proteins participate in the acclimation to both heat and cold. The significance of RNA processing for proper plant development is emphasized by the discovery that mutations in essential splicing genes frequently result in embryonic or post-embryonic lethality(14).

Auxin is important for establishing cell identity and polarity in embryos(15),(16) and in adult plants, it affects diverse processes such as root growth(17), leaf venation(18), and flower development(19). In addition, auxin biosynthesis, transport, perception, and signaling are known to modulate plant responses to different types of stress(20, 21). In particular, auxin biosynthesis and signaling have been implicated in thermomorphogenesis, the process in which plants elongate their hypocotyl and petioles, decrease leaf blade growth, and reposition their leaves to improve airflow for cooling during increased temperatures(22). A central regulator of ambient temperature signaling and growth is PHYTOCHROME INTERACTING FACTOR4 (PIF4)(22), which also plays important roles in the circadian clock(23), light signaling(24), and several phytohormone pathways, including auxin(25). PIF4 binds to multiple auxin-related genes, and its activity is essential for PIF4-mediated hypocotyl elongation during thermomorphogenesis(26). Cell expansion and hypocotyl elongation during thermomorphogenesis are, in part, controlled by several SMALL AUXIN UP RNA (SAUR) proteins, which regulate the proton ATPase activity in the plasma membrane and are themselves regulated by PIF4 and auxin(22).

Here, we show that the downregulation of *LSM7*, a core component of the spliceosome, results in increased seedling lethality, specifically at elevated temperatures (27°C). Transcriptomics and metabolomics analyses suggest that this hypersensitivity is due to an imbalance in auxin, most likely caused by improper mRNA processing of important auxin-responsive genes, including SAURs. In line with this hypothesis, constitutive overexpression of *SAUR19* partially rescued the temperature-sensitive phenotype of the hypomorphic *lsm7-2* mutant. Our study provides important insights into the mechanisms by which RNA processing modulates thermomorphogenesis through auxin homeostasis and signaling and suggests the possibility of modifying RNA processing to mitigate the effects of global warming on plant growth.

## Results

### *LSM7* is essential for embryo development

LSM7 is encoded by a single-copy gene (*AT2G03870*) and was previously shown to directly interact with PCP(27), which is indispensable for normal plant growth and development under cold and low ambient temperatures(12, 13). Yeast-two-hybrid analysis confirmed that LSM7 interacted with PCP and its paralog, PCP-like/SME2 **(Fig. S1a,b)**, and was therefore selected to further investigate its role in temperature-dependent plant development.

Insertion of a T-DNA in the third intron of *LSM7* (*lsm7-1*) **(Fig. 1a)** caused embryo lethality, with no homozygous mutants being recovered from a *lsm7-1* heterozygous (*lsm7-1*+/-) parent plant (>100 seeds tested). Furthermore, 27% of the ovules (n=139) in *lsm7-1*+/-plants were aborted, indicating that a single recessive mutation was causing the embryo lethality. 24% (31 out of 130) of the embryos analyzed in detail were arrested at the late globular stage **(Fig. 1b)**. In the same siliques, *lsm7-1*+/- and wildtype embryos in the non-aborted ovules had already reached the bend cotyledon stage **(Fig. 1b)**. *lsm7-1* embryos also displayed other defects, such as enlarged suspensor, hypophysis, and aberrant division patterns in the embryo **(Fig. 1b)**. Expression of *LSM7* coding (*pLSM7::cdsLSM7::tLSM7*) or genomic (*pLSM7::gLSM7::tLSM7*) sequence under its regulatory sequences rescued *lsm7-1* embryogenesis defects **(Fig. 1c)**, confirming that loss of *LSM7* was causing the embryo lethality in *lsm7-1*. Overall, our results identified LSM7 as an essential gene, complete loss-of-function of which resulted in early embryo abortion.

**Figure 1.**
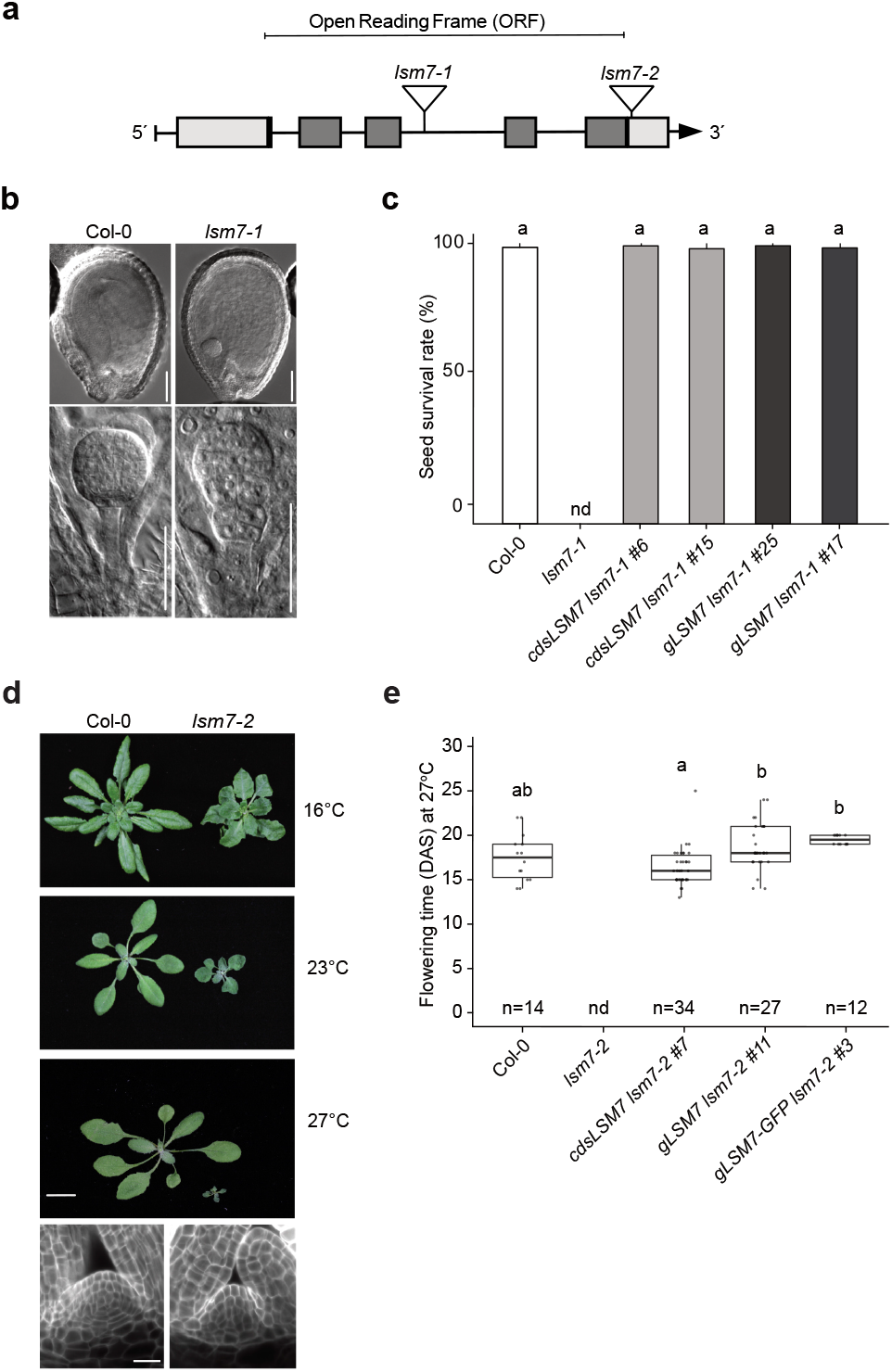
LSM7 is essential for embryo development and proper plant growth at low and high ambient temperatures. **a** Scheme of the *LSM7* gene locus and the T-DNA mutants used in this study. Exons are represented by grey boxes, UTRs are represented by light grey boxes, introns are represented by lines, and inverted triangles indicate T-DNA insertions. **b** Wildtype and *lsm7-1-/-* embryos collected from *lsm7-1+/-* plants. The scale bar is 100μm. **c** Seed survival rate (%) for Col-0 (white bar), T2 *pLSM7::cdsLSM7::tLSM7 lsm7-1* (light grey bar) and *pLSM7::gLSM7::tLSM7 lsm7-1* (dark grey bar) rescue lines. Error bars indicate standard deviation and letters significant differences for p<0.001 (t-test). **d** Wildtype and *lsm7-2* plants grew at different temperatures (the scale bar is 1cm), with an overview of the shoot apical meristem in seven-day-old wildtype and mutant seedlings (23°C). Cell organization is observed in median longitudinal confocal sections (the scale bar is 20μM). E, Epidermis; C, Cortex; En, Endodermis; St, Stele. **e** Ectopic expression of *LSM7* rescues *lsm7-2* lethality at 27°C. T1 plants expressing different constructs in *lsm7-2* background were scored at 27°C. One-way ANOVA was used for statistical analysis. Letters above the box plots indicate statistically different categories (p ≤0.05). Error bars are standard deviation (SD) of mean values. n, Number of individual plants; #, line numbering.

### An LSM7 hypomorphic mutant is temperature-sensitive

In contrast to *lsm7-1, lsm7-2* carries a T-DNA insertion in the 3’ UTR **(Fig. 1a)**, resulting in severe knock-down of *LSM7* expression **(Fig. S2a)**. This hypomorphic mutant is viable but displayed pleiotropic defects including a smaller rosette diameter, reduced leaf curvature, irregular leaf margins, darker color, and smaller shoot apical meristem **(Fig. 1d)**. Interestingly, these phenotypes increased in severity at higher temperatures **(Fig. 1d)**, and more than 80% of seedlings grown at 27°C eventually died. Expression of *LSM7* (*pLSM7::gLSM7::tLSM7* or *pLSM7::cdsLSM7::tLSM7*) fully rescued the *lsm7-*2 phenotype (>25 independent T1 lines) (**Fig. 1e**), confirming that the downregulation of *LSM7* is causal for the developmental defects in the mutant. Importantly, *LSM7* expression was significantly reduced in *lsm7-2* compared to wildtype (Col-0) at all three temperatures tested **(Fig. S2a)**, indicating that the differences in phenotype severity at the different temperatures were not due to differences in the degree of knock-down. Furthermore, LSM7 protein is ubiquitously expressed (*pLSM7::gLSM7-GFP::tLSM7*) in seven-day-old wildtype seedlings and neither its abundance nor subcellular localization was affected by temperature change **(Fig. S2b,c)** and can therefore not explain the changes in phenotype severity at elevated temperatures.

### *lsm7-2* has widespread consequences on the transcriptome

To better understand the molecular process that underlie the temperature-sensitive nature of the hypomorphic *lsm7-2* mutant we performed strand-specific RNA-Seq. Seedlings were initially grown at 23°C for nine days and then shifted to 16°C or 27°C or maintained at 23°C for a further 3h and 24h **(Fig. 2a)**. Principal component analysis revealed that 31.77% and 24.58% of the variance could be attributed to the different genotypes and temperature, respectively **(Fig. 2b)**. Differentially expressed (DE) and differentially alternatively spliced (DAS) genes were identified for each time point at the three temperatures by comparing the mutant and wildtype transcriptomes **(Fig. 2c; Dataset S1,2)**. Unsurprisingly, given the major effect of the genotype on the transcriptome, we found a similar number of genes up or downregulated at the three temperatures. Nevertheless, exposure to high and low temperatures increased the number of DE genes between *lsm7-2* and Col-0 **(Fig. 2d; Dataset S1)**. As expected for a mutant involved in mRNA processing, we observed more upregulated DAS genes in *lsm7-2* versus Col-0 in all conditions **(Dataset S2)**. GO term analysis of DE genes demonstrated the central role of LSM7 in controlling genes involved in key processes such as growth, development, hormone signaling, and responses to environmental changes (**Fig. 2e, Dataset S3**).

**Figure 2.**
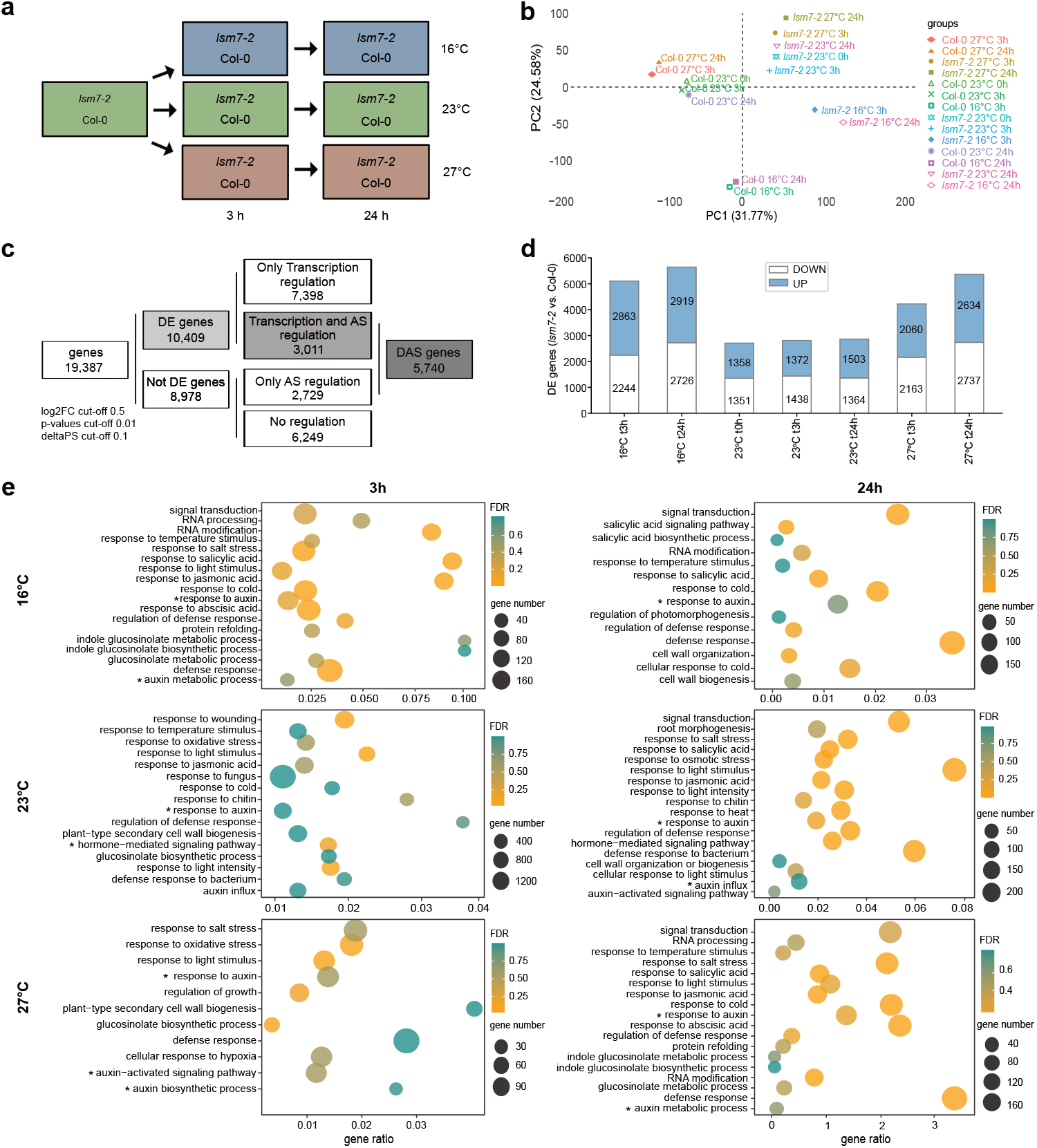
*lsm7-2* transcriptome upon shifts in ambient temperature. **a** RNA-seq experimental design. **b** Principal Component Analysis (PCA) for RNA-seq samples. Each data point from the RNA-seq data reflects the average gene expression (n = 3). **c** Number of genes or transcripts that are only regulated by transcription (DE), only by alternative splicing (AS), or both transcription and alternative splicing (DAS) are shown. **d** Number of differentially expressed genes (DEGs; log_2_FC *lsm7-2*/Col-0) in each tested condition detected by RNA-seq divided by up-or down-regulation. **e** Selected Biological Processes (BPs) for enriched Gene Ontologies (GOs) for DEGs of *lsm7-2* compared to Col-0 of nine-day-old seedlings after initial (3h) and late (24h) exposure to different ambient temperatures (16°C, 23°C and 27°C). The circle sizes and the false discovery rate (FDR) by colours represent the number of genes. Stars highlight the auxin-related GOs.

### LSM7 modulates auxin metabolism and homeostasis

Interestingly, GO categories related to hormone signaling pathways, especially auxin, were overrepresented in all experimental conditions **(Fig. 2e; Dataset S3**). DE and DAS auxin-related genes included many members of the *GRETCHEN HAGEN 3* (*GH3*), *AUXIN/INDOLE-3-ACETIC ACID* (*Aux/IAA*), *AUXIN RESPONSE FACTOR* (*ARF*), and *SMALL AUXIN UP-REGULATED RNA* (*SAUR*) families (**Dataset S2**). Importantly, even though many of these genes were already differentially regulated at 23°C, misregulation of these gene families increased at elevated or cooler ambient temperatures (**Dataset S2**), indicating that LSM7 plays a crucial role in regulating auxin homeostasis and signaling across the entire ambient temperature range (**Fig. 2e**).

In line with our transcriptome analyses, metabolic analysis revealed significantly elevated levels of the auxin precursors IAN and IAM in *lsm7-2* at all three temperatures **(Fig. S3a)**. However, free IAA was induced significantly only in mutants grown at 23°C **(Fig. S3b)**, suggesting plants strive to maintain IAA homeostasis, possibly by activating IAA catabolism. In agreement with this hypothesis, *GH3* genes encoding proteins involved in conjugating IAA to amino acids for storage or degradation were highly overrepresented among the DE and DAS genes in *lsm7-2* **(Fig. S4; Dataset S1,2)**. This induction was reflected in a significant increase of IAA-Asp, IAA-Glu, and IAA-Ala conjugates in *lsm7-2* at all three temperatures **(Fig. S3)**. Furthermore, other genes involved in IAA conjugation and catabolism, including *ILR1, DAO2*, and *UGT74D1*, were also significantly upregulated **(Fig. S4)**, correlating with an increase in oxIAA and oxIAA-Glc in *lsm7-2* **Fig. S3b)**. Interestingly, we observed increased IAA-Glc only after shift to 27°C, in agreement with a previous report^4^ **(Fig. S3b)**. Taken together our results indicate that the pool of auxin is increased in *lsm7-2*. The concomitant downregulation of the IAA influx genes and the upregulation of IAA efflux genes in *lsm7-2* **(Fig. S4)** likely reflects the plant’s effort to balance cellular auxin levels.

### LSM7 is required for hypocotyl elongation during thermomorphogenesis

The finding that the *lsm7-2* phenotype was strongly enhanced at 27°C even though our transcriptome and metabolome analyses detected misregulation of auxin homeostasis and signaling across the entire examined temperature range suggested that LSM7 might participate in regulating thermomorphogenesis. Supporting this hypothesis, we found that *PIF4*, a key regulator of thermomorphogenesis, was significantly downregulated in *lsm7-2* after 24h **(Fig. 3a)**. We, therefore, investigated if LSM7 could act through PIF4 and auxin to modulate thermomorphogenesis.

**Figure 3.**
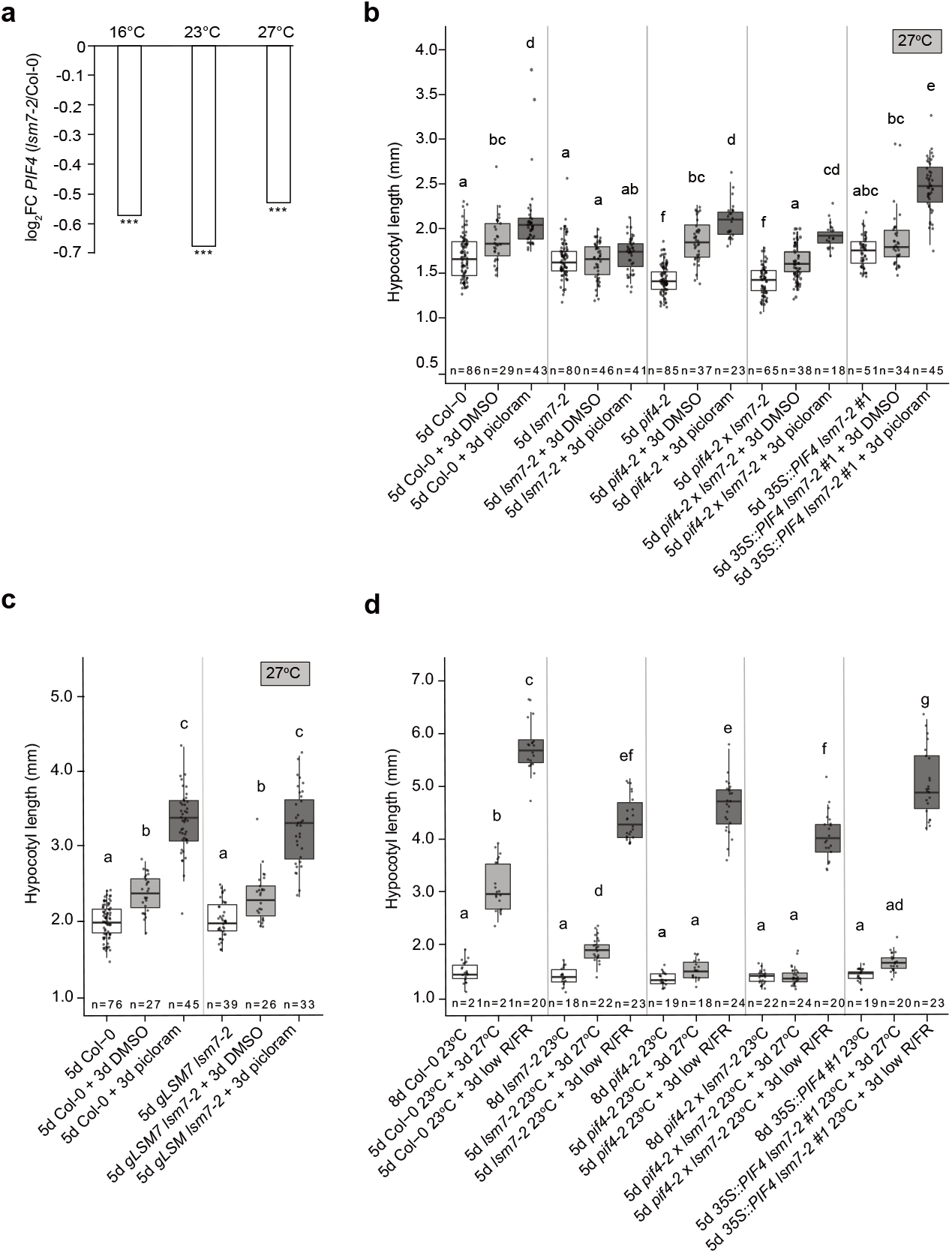
*lsm7-2* hypocotyl elongation defect is temperature-dependent. **a** RNA-seq expression of *PIF4* (log2FC *lsm7-2*/Col-0) in 10 days-old seedlings. The significance of FC values is indicated by three asterisks (padj<0.001). **b-d** Hypocotyl length of seedlings grown for five days at the indicated temperature and then transferred to the indicated treatment for three days or maintained at 23°C. The number of seedlings (n) is shown at the bottom of each box. Different letters denote statistical differences (p < 0.05) among samples as assessed by one-way ANOVA and Tukey HSD. Boxes indicate the first and third quartiles and the whiskers indicate the minimum and maximum values, the black lines within the boxes indicate the median values and grey dots mark the individual measurements. d, number of days. The different treatments are coloured with different shades of grey.

We first established that Col-0 hypocotyls elongated significantly over three days and that this effect was further enhanced in response to treatment with exogenous auxin (picloram) **(Fig. 3b)**. In contrast, hypocotyl length remained constant in *lsm7-2* and was significantly shorter than that of Col-0 by the end of the treatment **(Fig. 3b)**, indicating that *LSM7* was required for basal and auxin-induced hypocotyl elongation. We further observed that the hypocotyls of five-day-old *pif4-2* and *pif4-2 lsm7-2* plants were significantly shorter than those of Col-0, *lsm7-2*, and *35S::PIF4 lsm7-2* **(Fig. 3b)**, confirming that hypocotyl growth is in part mediated by PIF4. However, hypocotyls of both *pif4-2* and *pif4-2 lsm7-2* grew throughout the experiment and this growth was further enhanced by exogenous auxin. Interestingly, overexpression of *PIF4* (*35S::PIF4*) by itself did not promote hypocotyl elongation in *lsm7-*2 **(Fig. 3b)**, indicating that PIF4 requires *LSM7* to induce hypocotyl growth. However, the application of picloram induced strong hypocotyl elongation in *35S::PIF4 lsm7-2* **(Fig. 3b)**, demonstrating that exogenous auxin when combined with high levels of PIF4 was able to partially compensate for the reduced *LSM7* expression in *lsm7-2*. Supporting this notion, the response to exogenous auxin was fully restored in the *gLSM7 lsm7-2* rescue lines **(Fig. 3c)**. Surprisingly, picloram application also resulted in moderate hypocotyl elongation in *pif4-2* and *pif4-2 lsm7-2*, indicating that auxin-mediated hypocotyl elongation is not exclusively regulated by PIF4.

To test whether the function of LSM7 was specific to temperature we determined if hypocotyl elongation in *lsm-7-2* was also affected under shade-mimicking conditions (low red/far-red light ratio). We found that shade-mimicking conditions promoted hypocotyl elongation close to wildtype levels and that this effect was largely independent of PIF4 **(Fig. 3d)**. Taken together, our results suggest that the temperature sensitivity of *lsm7-2* can at least in part be explained by the failure to execute thermomorphogenesis responses such as hypocotyl elongation. Our results further indicate that PIF4 and LSM7-dependent auxin signaling act in parallel or converge on the same targets to achieve this effect.

### Overexpression of SAUR19 rescues the *lsm7-2* hypocotyl elongation defect

Since PIF4 activity is reduced in *lsm7-2* and treatment with picloram combined with PIF4 overexpression could restore *lsm7-2* hypocotyl defect **(Fig. 3b)**, we explored our transcriptome data for common PIF4 and auxin targets that could explain the *lsm7-2* hypocotyl elongation defects. Among the auxin-responsive genes that were misregulated in our transcriptome analyses **(Fig. 2e, Dataset S1)**, the *SAUR* family, key regulators of auxin-, temperature-, and PIF4-mediated growth responses such as hypocotyl elongation during thermomorphogenesis and shade avoidance(28– 30), was the most severely affected. Of the 79 *Arabidopsis SAUR* genes, 42 were DE, of which 35 were significantly downregulated in *lsm7-2* **(Fig. 4a)**. Of these, *SAUR19* was the most strongly downregulated *SAUR* gene at elevated ambient temperature. Importantly, *SAUR19* overexpression resulted in full rescue of *lsm7-2* hypocotyl elongation **(Fig. 4b)**, indicating that restoration of *SAUR19* expression could partially relieve developmental defects of *lsm7-2*.

**Figure 4.**
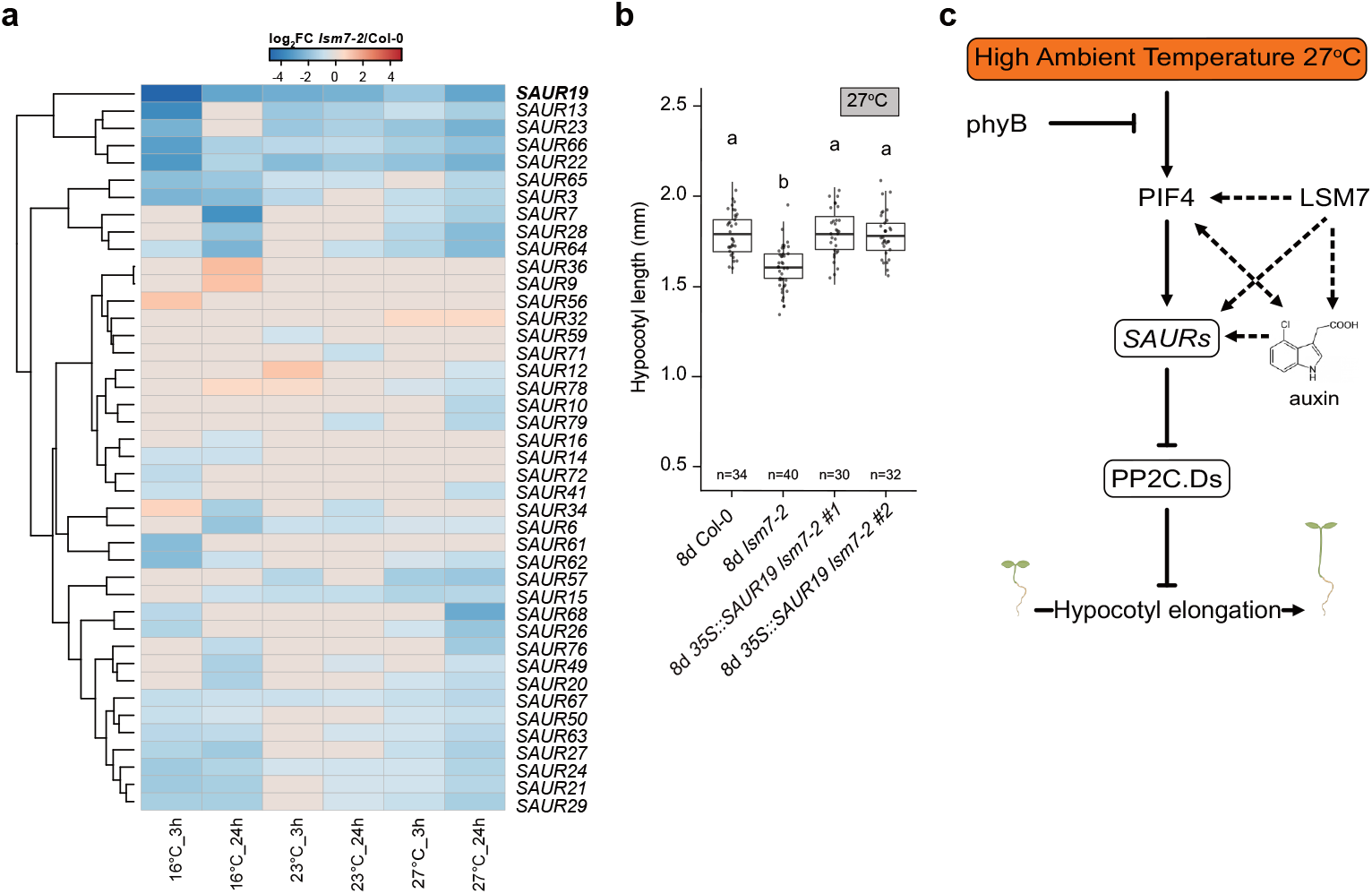
*SAUR* genes are misregulated in *lsm7-2*. **a** Heatmaps of significant differentially expressed *SAUR* genes (log2FC *lsm7-2*/Col-0) in seedlings exposed to different ambient temperatures, organized according to their function. Shades of red indicate up-regulated genes, and shades of blue indicate downregulated genes in *lsm7-2*. **b** Hypocotyl length of 8-day-old seedlings grown at 27°C. The number of seedlings (n) is shown at the bottom of each box. Different letters denote statistical differences (p < 0.05) among samples as assessed by one-way ANOVA and Tukey HSD. Boxes indicate the first and third quartiles and the whiskers indicate the minimum and maximum values, the black lines within the boxes indicate the median values and grey dots mark the individual measurements. **c** Working model summarizing our results on the LSM7 role at high ambient temperatures.

Overall, our findings highlight the role of RNA processing in general and LSM7 in particular in temperature and hormonal signaling. LSM7 contributes to auxin homeostasis and signaling by controlling the transcriptional efficiency and accuracy of essential temperature and auxin-responsive genes. Ultimately, the integration of PIF4 and LSM7-mediated modulation of auxin-responsive genes such as the *SAURs* controls hypocotyl elongation at high ambient temperature, a key aspect of thermomorphogenesis and an important developmental response at elevated temperatures (**Fig. 4c)**.

## Discussion

Recent studies have shown that splicing-associated proteins are essential for plant growth and development under suboptimal temperatures(7, 10, 12, 13). Here, we found that LSM7, a mRNA processing factor involved in RNA splicing and the NMD pathway, is essential for *Arabidopsis* embryo and plant development **(Fig. 1c)**. In contrast to the embryo lethal *lsm7-1* allele, the knock-down mutant *lsm7-2* is viable, but displays pleiotropic developmental defects that become more severe with warmer temperatures, culminating in seedling lethality at temperatures as low as 27°C **(Fig 1e)**. Although other Sm and LSm proteins have been implicated in temperature responses(7), *lsm7-2* is the only mutant with reduced viability under such moderately warm ambient temperatures. This unique feature distinguishes LSM7 from other mRNA processing factors and highlights its critical role in temperature acclimation. Importantly, temperature is not regulating *LSM7* gene expression or protein *per se* **(Fig. S2)**, suggesting that the temperature sensitivity of *lsm7-2* is due to its function in processing mRNAs of important downstream genes.

Transcriptomic analyses **(Fig. 2)** verified that downregulation o*f LSM7* modulates the expression of genes involved in several key biological processes **(Fig. 2e)**, including auxin homeostasis (biosynthesis, conjugation, and degradation) and signaling. Auxin regulates plant growth and development in a concentration and tissue-dependent manner and controls fundamental processes such as cell division, expansion, and differentiation(31). In agreement with the transcriptome data, targeted auxin-associated metabolite analysis **(Fig. S4)** showed that levels of free auxin (IAA), the precursors IAM and IAN, and auxin conjugates such as IAA-Ala, IAA-Glu, and IAA-Asp accumulate in *lsm7-2*. The latter also correlates with the upregulation of the *GH3s* genes **(Fig. S3)**, which catalyzes the conjugation of IAA with amino acids for inactivation and storage(32). The finding that products of IAA oxidation, such as oxIAA and oxIAA-Glc, are significantly accumulated in *lsm7-2* supports our hypothesis that downregulation o*f LSM7* promotes IAA storage and inactivation to prevent overaccumulation of bioactive IAA(33). The misregulation of genes involved in IAA influx and efflux, respectively, are also consistent with this hypothesis **(Fig. S3)**.

Despite these compensatory mechanisms, auxin homeostasis and expression of key auxin signaling genes such as *AUX/*IAAs and *ARFs* are perturbed in *lsm7-2*. It is known that warm temperature promotes hypocotyl growth via the PIF4-auxin pathway, which regulates the expression of auxin-responsive genes required for elongation(24). Unexpectedly, the application of auxin was not sufficient to restore the hypocotyl elongation defect in *lsm7-2* at high ambient temperature, whereas *PIF4* overexpression had a minor effect **(Fig. 3b)**, suggesting that LSM7 could act in parallel to the PIF4-auxin pathway to converge on a common set of targets. In agreement with this hypothesis, *SAURs* were among the most misregulated gene families **(Fig. 4a)**. SAURs are involved not only in thermomorphogenesis(28) but also in other processes, such as in leaf senescence, or cell division and expansion(34). Their broad range of functions could not only explain the hypocotyl defect in high ambient temperatures but also contribute to the developmental defects of *lsm7-2* in normal ambient temperatures. *SAUR* genes are rapidly induced by both auxin and PIF4. The SAUR proteins, in turn, interact with PP2C.D enzymes to inhibit their repressive activity on H+-ATPase, which reduces the apoplastic pH and promotes cell elongation by cell wall loosening and increasing water uptake(35). SAURs are divided into groups based on their partially overlapping functions. The group of *SAUR19–24* has previously been identified as downstream targets of the PIF4-auxin pathway(28). Our transcriptomic analyses ranked *SAUR19* as the most downregulated *SAUR* gene and overexpression of *SAUR19* could restore hypocotyl elongation in *lsm7-2* at 27°C **(Fig. 4b)**. Taken together, our results identify LSM7 as an essential factor for auxin-responsive gene transcription at high ambient temperature to ensure successful integration of PIF4 and auxin signaling. Importantly, the effect of LSM7 on hypocotyl elongation seems to be specific to temperature as shade-mimicking conditions, which also trigger the elongation of hypocotyls via the PIF-auxin pathway by inducing *SAURs* and other auxin-responsive genes(24, 36, 37), could restore hypocotyl elongation in *lsm7-2* close to wildtype levels **(Fig. 3d)**. A possible explanation for these differences could be that warm temperatures are known to increase the expression of auxin-responsive genes moderately, whereas low R/FR ratios have a much more pronounced effect(38). Notwithstanding, this will be an interesting aspect to further investigate in future work.

Taken together, our study provides valuable insights into the role of a basal component of the splicing and RNA degradation machinery, LSM7, in responding to small changes in ambient temperature. We demonstrated that LSM7 is a key regulator required for efficient mRNA expression and processing of numerous auxin-related genes, from auxin biosynthesis to signaling and transport, thereby coordinating the response of plants to ambient temperature. Based on these findings, we propose that the misregulation of auxin genes, and especially the downregulation of numerous *SAUR* genes, prevents *lsm7-2* from correctly executing thermomorphogenesis, ultimately leading to increased seedling lethality at moderately warm temperatures.

## Materials and Methods

### Plant material and growth conditions

*lsm7-1* (SALK_066076C), *lsm7-2* (SALK_065217), and *pif4-2* (SAIL_1288_E07) seeds were obtained from Nottingham Arabidopsis Stock Center (NASC). Rescue lines (c*LSM7*, g*LSM7* or *gLSM7-GFP*) were created using the GreenGate system(39). The promoter (2140 bp upstream of the start codon) and terminator (256 bp downstream of the stop codon) regions of the *LSM7* gene (g*LSM7*) were amplified from Col-0 seedlings and cloned in pGGA000 (p*LSM7*), and pGGE000 (t*LSM7*), respectively. The c*LSM7* was also amplified from Col-0 cDNA and cloned in the pGGC000 and the pGGF004 for BASTA resistance as a plant selection marker. The pGGZ003 was used as the destination vector for all the final constructs. Col-0, *lsm7-1* or *lsm7-2* plants were grown at 23°C and transformed by floral dipping using *Agrobacterium tumefaciens*-mediated gene transfer(40). Overexpression lines for *PIF4* (*35S::PIF4*) and *SAUR19* (*35S::SAUR19)* were created by transforming the *lsm7-2* mutant. The *35S::SAUR19* lines were created by amplifying *SAUR19* CDS from Col-0 cDNA and cloned in the pGGC000 and the pGGF004 for BASTA resistance as a plant selection marker, in final GreenGate vector pGGZ003. The plasmid pCF402 (*35S::PIF4-HA*) to create the *35S::PIF4* lines was a gift from Prof. Christian Fankhauser(41). Transformants were selected by BASTA application. or GFP-seed coat screening for the overexpression lines *35S::PIF4* **(Dataset S4)**.

Seeds were surface sterilized with 70% ethanol and then stratified at 4°C in the dark for 72 hours before sowing on soil or half-strength MS medium plates. Plants were cultivated in long-day conditions (16 hours light/8 hours dark) in Percival chambers with full-range LED illumination and photosynthetically active radiation (PAR) of 120-150 µmol^m-2s-1^, controlled humidity (RH 70%) at three ambient temperatures (16°C, 23°C, and 27°C) as specified in the text. Flowering time was recorded in soil-growth seedlings when apical meristems were visible (DAS – days after sowing), and rosette and cauline leaves were then counted. Hypocotyl elongation measurements were taken in plate-growth seedlings. Briefly, seedlings were grown on vertical plates in Percival chambers at 23°C for five days under light and grow conditions as described above. High ambient temperature (27°C) and low R/FR treatment started on day five at ZT6. For picloram (Sigma-Aldrich, Steinheim, Germany, P5575) treatment, nylon meshes were transferred on day five before the temperature shift to half-strength MS medium with 2µM picloram and 0.1% dimethyl sulphoxide (DMSO) for mock treatment. Seedling imaging and measurements were performed on day eight with the Fiji package from ImageJ software (https://fiji.sc).

### Seed survival determination

For seed survival determination, siliques at mature stages of development were dissected under a Leica MZ16 stereomicroscope, and seeds from 20 siliques of 15 individual plants were analyzed. The percentage represents the developed versus the total number of seeds counted in the siliques for each line. Plants were grown at 23°C and long day conditions as described above.

### Microscopic analysis of embryos

For microscopic analysis, immature seeds were dissected by hand and cleared in Hoyer’s solution (7.5g gum arabic, 5 mL glycerol, 100 g chloral hydrate and 30 mL water, diluted 2:1 with 10% (w/v) gum arabic solution) as described previously(42). Three days after incubation, differential interference contrast (DIC) images were taken with a Zeiss Axio Imager equipped with AxioCam HRc camera.

### Protein-protein interactions

The coding sequence of *PCP* (*AT2G18740*), *PCP-like* (*AT4G30330*), *LSM5* (*AT5G48870*), *LSM7* (*AT2G03870*) and its truncated form *LSM7-1t* were cloned into the modified yeast vectors pGADT7 or pGBKT7(43) **(Dataset S4)**. Yeast two-hybrid (Y2H) assays were conducted, and plasmids were co-transformed in AH109 yeast strain. Transformants and interactions were screened on SD-glucose medium lacking leucine (-L), tryptophan (-W), and histidine (-H). Auto-activation was determined for all bait vectors to exclude false positive interactions.

### LSM7 subcellular localization and quantification

For LSM7 protein subcellular quantification, *lsm7-2* and Col-0 seedlings bearing the *pLSM7:GFP:gLSM7:tLSM7* construct were grown for nine days at 23°C in LD conditions and shifted for 24h to 16°C or 27°C, or kept at 23°C before been live imaged. To determine GFP-LSM7 enrichment in the nucleus, ten-day-old rescue seedlings were counterstained in ClearSee with 10 µg/mL Hoechst 33342 for one day(44). To minimize the risk of exposing the cells to any other factor than the temperature treatment, gLSM7-GFP observations were performed in seedlings mounted in the water right after taking them out of incubators. Imaging of epidermal cells in the root tip was recorded in z-plane stacks. For measurements, optical stack slices in a 5µM-thick section, spaced 1µM between each other, were converted into a maximum projection file in 8-bit depth. Mean intensity was quantified in regions of interest (ROI) in the nucleus and cytoplasm ranging between 1 and 2 µm^2^. The mean intensity, which is the average fluorescence per area unit, was calculated by adding the fluorescence intensity (in an 8-bit scale of 0 to 255) of each pixel in an area, divided by the number of pixels in that area. At least two cells showing clearly visible nuclei were considered per stack file, with each cell associated with an ROI in the nucleus and another in the cytoplasm. The mean intensities of measurements of different pictures were analyzed together. Images were processed and analyzed using the Fiji package of ImageJ (https://fiji.sc).

### LSM7 protein expression analysis

For LSM7 protein expression, nine-day-old *lsm7-2* and Col-0 seedlings bearing the *pLSM7:GFP:gLSM7:tLSM7* construct were grown at 23°C in LD conditions and shifted to 16°C or 27°C, or kept at 23°C for 24h. Samples were collected at ZT6, and total proteins were extracted using the TCA/Acetone method. Proteins were resolved in 4-15% pre-cast SDS-PAGE (Biorad cat nº 4561085) at 80V for two hours and transferred to PVDF membrane (Immobilon-P; Millipore) for immunoblot analysis. Two identical gels and relative membranes were made at the same time for the detection of GFP-LSM7 and tubulin (loading control). First, the membranes were blocked with 3% BSA in PBS for two hours. Then, one membrane was incubated with anti-GFP antibody (1/4000 Abcam cat. nº Ab290) with 3% BSA in PBS and the other with anti-Tubulin (1/10000 Agrisera cat. nº AS10-680) in 3% skimmed milk in PBS for two hours at room temperature. Both membranes were washed with TBS-T (Tween 20; 50-mM Tris, 150-mM NaCl, and 0.05% [v/v] Tween 20) three times and incubated with the anti-rabbit IgG-HRP antibody (1/20,000, Agrisera cat. nº AS10-1014) in 3% w/v BSA (anti-GFP) or 3% milk (anti-tubulin) in PBS for two hours. After three washes with TBS-T, the membranes were incubated with Chemiluminescent Substrate (Amersham cat. nº RPN2232) for 1 min. Proteins were visualized with the Azure Imaging System (Azure Biosystems, Inc).

### Transcriptomic analysis

To analyze transcriptomic changes in response to different ambient temperatures, *lsm7-2* and Col-0 seedlings were grown for nine days at 23°C in LD conditions and shifted at ZT6 (time-point 0h) to 16°C, 27°C or kept at 23°C in LD and sampled after 3h and 24h. Three pools of ten seedlings’ aerial parts were collected per tested condition. Frozen samples were ground to a fine powder, and total RNA was extracted with Qiagen Plant RNeasy kit according to the manufacturer’s instructions and treated with DNAseI (Thermo Scientific). RNA concentrations and integrity were determined using Qubit RNA kit (Thermo Scientific) and Bioanalyser RNA Nano kit (Agilent). Strand-specific mRNA-Seq was conducted by Novogene using NEB Next® Ultra RNA Library Prep Kit for Illumina. Briefly, mRNA was purified from total RNA using poly-T oligo-attached magnetic beads. After fragmentation, the first-strand cDNA was synthesized using random hexamer primers, followed by the second-strand cDNA synthesis, followed by repair, A-tailing, adapter ligation, and size selection. After amplification and purification, the insert size of the library was validated on an Agilent 2100 and quantified using quantitative PCR (qPCR). Libraries were then sequenced on Illumina NovaSeq 6000 S4 flowcell with PE150. The data analysis was conducted using R and Bioconductor with some modifications. The quality of the sequences was confirmed using Trimmomatic(45) and FastQC. Reads were mapped to the AtRTD2-QUASI transcriptome annotation(46) using Salmon(47). After normalization, the profiles of the samples were assessed by principal component analysis (PCA). Differentially expressed (DE) genes between temperatures and genotypes across the time points were identified at each time point using limma-voom likelihood ratio tests after negative binomial fittings using the package in the 3D pipeline(48). Genes with False Discovery Rate (FDR)-corrected-values ≤ 0.05 and fold-change (log2) threshold of 0.5 were classified as differentially expressed **(Dataset S1,2)**. To identify processes potentially involved in the plant response to the different temperatures at the different time points, GO Enrichment Analysis was performed using online David software version 2021(49) **(Dataset S3)**.

### Gene expression analysis by quantitative real-time PCR (qPCR)

RNA samples were treated with DNaseI, RNase-free (Thermo Scientific), and then retro-transcribed using the RevertAid First Strand cDNA Synthesis kit (Thermo Fisher) according to the manufacturer’s instructions. Primers were designed using Primer 3 software^13^ with the criterion of generating amplified products from 80 to 180 bp with a Tm of 60 ± 1°C **(Dataset S4)**. The cDNA samples were used in qPCR reactions with LightCycler Sybr (Roche), using the CFX96 Real-time System (Biorad). A normalized expression ratio was calculated using an efficiency (E)-calibrated model, and the experimental significance was estimated through 2000 randomizations of Cq data in each experimental comparison using REST software (version mcs)^14^ Three technical and three biological replicates were performed for each sample.

### IAA and IAA-conjugate measurements

Col-0 and *lsm7-2* seedlings were grown for nine days at 23°C in LD conditions and shifted to ambient temperatures of 16°C or 27°C, or kept at 23°C for 24 hours. At ZT6, five pools of 20mg seedling shoots for each genotype at each temperature were collected in liquid nitrogen. The extraction, purification, and LC-MS analysis of endogenous IAA, its precursors, and metabolites were carried out according to Novák et al^15^. A bead mill (27 Hz, 10 min, 4°C; MixerMill, Retsch GmbH, Haan, Germany) was used to homogenize 20 mg of frozen material per sample, which was then extracted in 1 ml of 50 mM sodium phosphate buffer containing 1% sodium diethyldithiocarbamate and a mixture of 13C6-or deuterium-labeled internal standards. After centrifugation (14 000 RPM, 15 min, 4°C), the supernatant was divided into two aliquots. The first aliquot was derivatized using cysteamine (0.25 M; pH 8; 1 h; room temperature; Sigma-Aldrich); the second aliquot was immediately further processed as follows: the pH of the sample was adjusted to 2.5 by 1 M HCl and applied on a preconditioned solid-phase extraction column, Oasis HLB (30 mg 1 cc, Waters Inc., Milford, MA, USA). After sample application, the column was rinsed with 2 ml of 5% methanol. The compounds of interest were then eluted with 2 ml of 80% methanol. The derivatized fraction was similarly purified. Mass spectrometry analysis and quantification were performed by an LC-MS/MS system comprising a 1290 Infinity Binary LC System coupled to a 6490 Triple Quad LC/MS System with Jet Stream and Dual Ion Funnel technologies (Agilent Technologies, Santa Clara, CA, USA).

## Statistical analysis

Each section provides information on statistical tests, which are included in figures and figure legends (P-value, P-value levels, and sample number).

## Data availability

The RNA-seq data generated in this study have been deposited at the European Nucleotide Archive (ENA, https://www.ebi.ac.uk/ena) under the accession number PRJEB77534. Lists of differentially expressed genes and transcripts are made available as Supplementary Files. Other data supporting this study’s findings are available from the corresponding author upon request.

## Acknowledgments

The authors acknowledge the facilities and technical assistance of the Umeå Plant Science Center (UPSC). More in particular, we would like to thank Dr Nicholas Delhome and the UPSC Bioinformatics facility (UPSCb, Umeå, Sweden, https://www.upsc.se/platforms/upsc-bioinformatics-facility.html) for the bioinformatics support; the personnel working at the Microscopy facility (https://www.upsc.se/platforms/microscopy-facility.html); Dr Jan Šimura and the Swedish Metabolomics Center (SMC, Umeå, Sweden, www.swedishmetabolomicscentre.se) for carrying out the auxin metabolites’ analysis; and all members of the MS group for the scientific discussions and comments on the text. Figure 4 was partially created with a fully licensed Biorender.com. The computational data analysis was enabled by resources in projects NAISS 2024/22-1054 and NAISS 2024/23-466 to SMN, provided by the National Academic Infrastructure for Supercomputing in Sweden (NAISS) at UPPMAX, funded by the Swedish Research Council through grant agreement no. 2022-06725. This work was supported by the German Research Foundation (Deutsche Forschungsgemeinschaft -DFG) (BA3356/3-1, BA3356/4-1, and SFB1101/B12) to MB and the Knut and Alice Wallenberg Foundation (KAW 2018.0202) and FORMAS (2023-01077) to MS.

## Notes

### Competing Interest Statement

The authors have declared no competing interest.

### Summary of Updates

Additional experiments were conducted to investigate the role of LSM7 in thermomorphogenesis, particularly in relation to hypocotyl elongation in response to warm ambient temperatures. Furthermore, we demonstrate that the constitutive expression of SAUR19 partially rescues the hypocotyl elongation defects associated with the hypomorphic lsm7-2 allele. The revised manuscript provides new insights into the molecular mechanism underlying the temperature-sensitive phenotype of lsm7-2, which was not addressed in the original submission.

## References

1. C. P. G. Calixto, et al., Rapid and Dynamic Alternative Splicing Impacts the Arabidopsis Cold Response Transcriptome. Plant Cell 30, 1424–1444 (2018).

2. N. V. Lekontseva, E. A. Stolboushkina, A. D. Nikulin, Diversity of LSM Family Proteins: Similarities and Differences. Biochemistry 86, S38–S49 (2021).

3. W. He, R. Parker, Functions of Lsm proteins in mRNA degradation and splicing. Curr. Opin. Cell Biol. 12, 346–350 (2000).

4. D. Wu, et al., Lsm2 and Lsm3 bridge the interaction of the Lsm1-7 complex with Pat1 for decapping activation. Cell Res. 24, 233–246 (2014).

5. M. Arribas-Layton, D. Wu, J. Lykke-Andersen, H. Song, Structural and functional control of the eukaryotic mRNA decapping machinery. Biochim. Biophys. Acta 1829, 580–589 (2013).

6. T. Kurosaki, M. W. Popp, L. E. Maquat, Quality and quantity control of gene expression by nonsense-mediated mRNA decay. Nat. Rev. Mol. Cell Biol. 20, 406–420 (2019).

7. V. Dikaya, et al., Insights into the role of alternative splicing in plant temperature response. J. Exp. Bot. (2021). 10.1093/jxb/erab234.

8. J. L. Mateos, et al., PICLN modulates alternative splicing and light/temperature responses in plants. Plant Physiol. 191, 1036–1051 (2023).

9. C. Perea-Resa, T. Hernández-Verdeja, R. López-Cobollo, M. del Mar Castellano, J. Salinas, LSM proteins provide accurate splicing and decay of selected transcripts to ensure normal Arabidopsis development. Plant Cell 24, 4930–4947 (2012).

10. M. Okamoto, et al., Sm-Like Protein-Mediated RNA Metabolism Is Required for Heat Stress Tolerance in Arabidopsis. Front. Plant Sci. 7, 1079 (2016).

11. C. Carrasco-López, et al., Environment-dependent regulation of spliceosome activity by the LSM2-8 complex in Arabidopsis. Nucleic Acids Res. 45, 7416–7431 (2017).

12. R. Huertas, et al., Arabidopsis SME1 Regulates Plant Development and Response to Abiotic Stress by Determining Spliceosome Activity Specificity. Plant Cell 31, 537–554 (2019).

13. G. Capovilla, et al., PORCUPINE regulates development in response to temperature through alternative splicing. Nat Plants 4, 534–539 (2018).

14. D. W. Meinke, Genome-wide identification of EMBRYO-DEFECTIVE (EMB) genes required for growth and development in Arabidopsis. New Phytol. 226, 306–325 (2020).

15. S. Verma, V. P. S. Attuluri, H. S. Robert, An Essential Function for Auxin in Embryo Development. Cold Spring Harb. Perspect. Biol. 13 (2021).

16. J. Friml, et al., Efflux-dependent auxin gradients establish the apical-basal axis of Arabidopsis. Nature 426, 147–153 (2003).

17. Y. Hu, et al., Cell kinetics of auxin transport and activity in Arabidopsis root growth and skewing. Nat. Commun. 12, 1657 (2021).

18. E. Mazur, I. Kulik, J. Hajný, J. Friml, Auxin canalization and vascular tissue formation by TIR1/AFB-mediated auxin signaling in Arabidopsis. New Phytol. 226, 1375–1383 (2020).

19. R. Aloni, E. Aloni, M. Langhans, C. I. Ullrich, Role of auxin in regulating Arabidopsis flower development. Planta 223, 315–328 (2006).

20. K. Kazan, Auxin and the integration of environmental signals into plant root development. Ann. Bot. 112, 1655–1665 (2013).

21. K. Shibasaki, M. Uemura, S. Tsurumi, A. Rahman, Auxin response in Arabidopsis under cold stress: underlying molecular mechanisms. Plant Cell 21, 3823–3838 (2009).

22. M. Quint, et al., Molecular and genetic control of plant thermomorphogenesis. Nat Plants 2, 15190 (2016).

23. Y. Nomoto, S. Kubozono, T. Yamashino, N. Nakamichi, T. Mizuno, Circadian clock- and PIF4-controlled plant growth: a coincidence mechanism directly integrates a hormone signaling network into the photoperiodic control of plant architectures in Arabidopsis thaliana. Plant Cell Physiol. 53, 1950–1964 (2012).

24. M. Legris, C. Nieto, R. Sellaro, S. Prat, J. J. Casal, Perception and signalling of light and temperature cues in plants. Plant J. 90, 683–697 (2017).

25. J. A. Stavang, et al., Hormonal regulation of temperature-induced growth in Arabidopsis. Plant J. 60, 589–601 (2009).

26. W. M. Gray, A. Ostin, G. Sandberg, C. P. Romano, M. Estelle, High temperature promotes auxin-mediated hypocotyl elongation in Arabidopsis. Proc. Natl. Acad. Sci. U. S. A. 95, 7197–7202 (1998).

27. S. Orchard, et al., The MIntAct project--IntAct as a common curation platform for 11 molecular interaction databases. Nucleic Acids Res. 42, D358–63 (2014).

28. K. A. Franklin, et al., Phytochrome-interacting factor 4 (PIF4) regulates auxin biosynthesis at high temperature. Proc. Natl. Acad. Sci. U. S. A. 108, 20231–20235 (2011).

29. H. van Mourik, A. D. J. van Dijk, N. Stortenbeker, G. C. Angenent, M. Bemer, Divergent regulation of Arabidopsis SAUR genes: a focus on the SAUR10-clade. BMC Plant Biol. 17, 245 (2017).

30. M. Bemer, et al., FRUITFULL controls SAUR10 expression and regulates Arabidopsis growth and architecture. J. Exp. Bot. 68, 3391–3403 (2017).

31. A. W. Woodward, B. Bartel, Auxin: regulation, action, and interaction. Ann. Bot. 95, 707–735 (2005).

32. K.-I. Hayashi, et al., The main oxidative inactivation pathway of the plant hormone auxin. Nat. Commun. 12, 6752 (2021).

33. J. M. Jez, Connecting primary and specialized metabolism: Amino acid conjugation of phytohormones by GRETCHEN HAGEN 3 (GH3) acyl acid amido synthetases. Curr. Opin. Plant Biol. 66, 102194 (2022).

34. N. Stortenbeker, M. Bemer, The SAUR gene family: the plant’s toolbox for adaptation of growth and development. J. Exp. Bot. 70, 17–27 (2019).

35. A. K. Spartz, et al., SAUR Inhibition of PP2C-D Phosphatases Activates Plasma Membrane H+-ATPases to Promote Cell Expansion in Arabidopsis. Plant Cell 26, 2129–2142 (2014).

36. J.-H. Jung, et al., Phytochromes function as thermosensors in Arabidopsis. Science 354, 886–889 (2016).

37. M. Legris, et al., Phytochrome B integrates light and temperature signals in Arabidopsis. Science 354, 897–900 (2016).

38. L. Bianchimano, M. B. De Luca, M. B. Borniego, M. J. Iglesias, J. J. Casal, Temperature regulation of auxin-related gene expression and its implications for plant growth. J. Exp. Bot. (2023). 10.1093/jxb/erad265.

39. A. Lampropoulos, et al., GreenGate---a novel, versatile, and efficient cloning system for plant transgenesis. PLoS One 8, e83043 (2013).

40. S. J. Clough, A. F. Bent, Floral dip: a simplified method for Agrobacterium-mediated transformation of Arabidopsis thaliana. Plant J. 16, 735–743 (1998).

41. S. Lorrain, T. Allen, P. D. Duek, G. C. Whitelam, C. Fankhauser, Phytochrome-mediated inhibition of shade avoidance involves degradation of growth-promoting bHLH transcription factors. Plant J. 53, 312–323 (2008).

42. M. Bayer, et al., Paternal control of embryonic patterning in Arabidopsis thaliana. Science 323, 1485–1488 (2009).

43. V. Zacharaki, et al., Impaired KIN10 function restores developmental defects in the Arabidopsis trehalose 6-phosphate synthase1 (tps1) mutant. New Phytol. 235, 220–233 (2022).

44. D. Kurihara, Y. Mizuta, Y. Sato, T. Higashiyama, ClearSee: a rapid optical clearing reagent for whole-plant fluorescence imaging. Development 142, 4168–4179 (2015).

45. A. M. Bolger, M. Lohse, B. Usadel, Trimmomatic: A flexible trimmer for Illumina sequence data. Bioinformatics 30, 2114–2120 (2014).

46. R. Zhang, et al., A high quality Arabidopsis transcriptome for accurate transcript-level analysis of alternative splicing. Nucleic Acids Res. 45, 5061–5073 (2017).

47. R. Patro, G. Duggal, M. I. Love, R. A. Irizarry, C. Kingsford, Salmon provides fast and bias-aware quantification of transcript expression. Nat. Methods 14, 417–419 (2017).

48. W. Guo, et al., 3D RNA-seq: a powerful and flexible tool for rapid and accurate differential expression and alternative splicing analysis of RNA-seq data for biologists. RNA Biol. 1–14 (2020). 10.1080/15476286.2020.1858253.

49. D. W. Huang, B. T. Sherman, R. A. Lempicki, Systematic and integrative analysis of large gene lists using DAVID bioinformatics resources. Nat. Protoc. 4, 44–57 (2009).

