## Supporting Figures 1-4 for "*Arabidopsis thaliana* LSM7 is essential for auxin-mediated regulation of *SAUR* genes and thermomorphogenesis"

**This PDF file includes:**

Figures S1 to S4
Legends for Datasets S1 to S4

**Other supporting materials for this manuscript include the following:**

Datasets S1 to S4

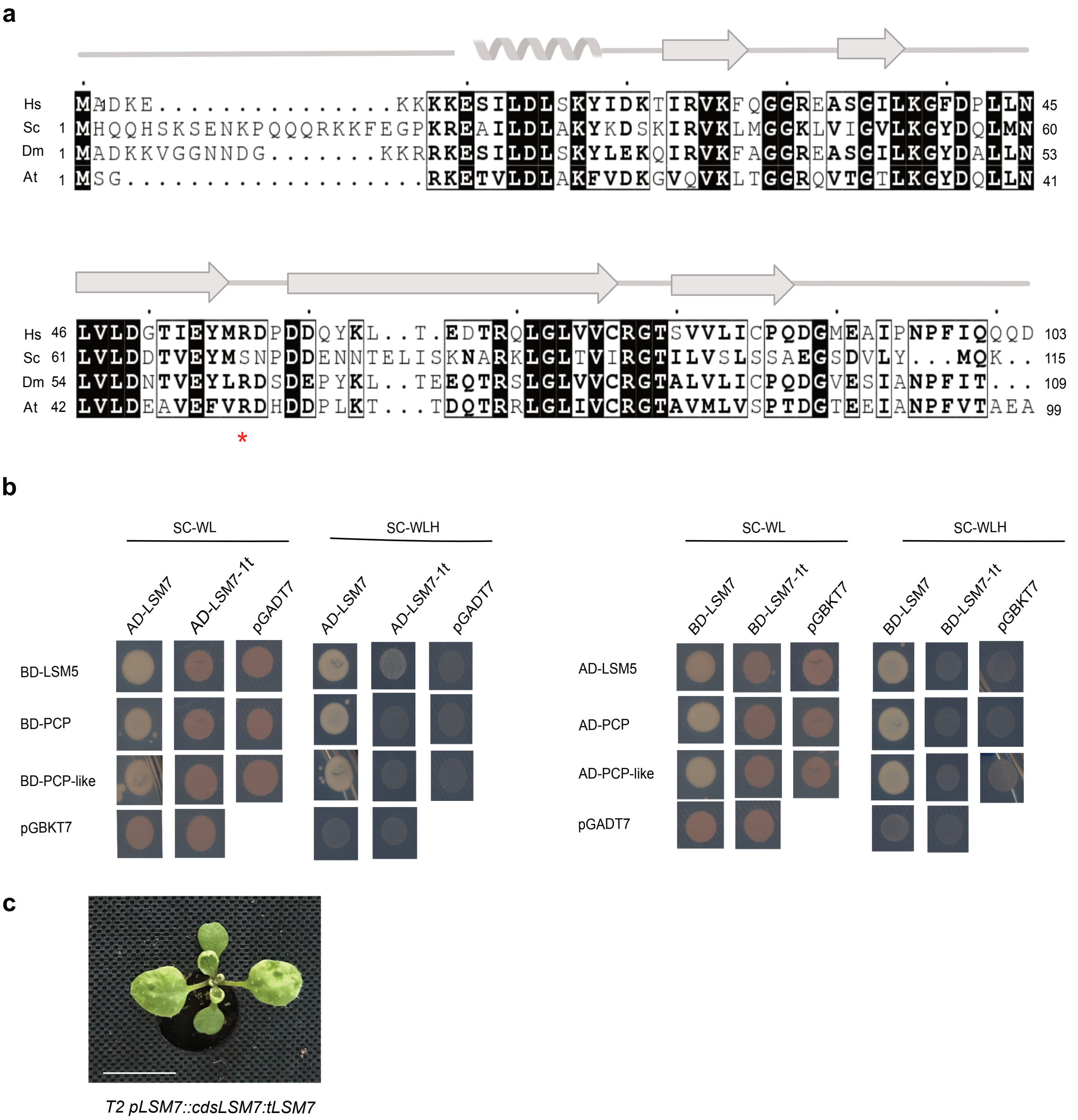


**Fig. S1. Interactome of LSM7 and SM/LSM selected partners. a** Schematic representation of LSM7 in different species. Above the amino acid sequence, the position of the helix and beta-sheets are reported. The red star indicates the end of the truncated protein in *lsm7-1*. Hs, *Homo sapiens*; Sc, *Saccharomyces cerevisiae*; Dm, *Drosophila melanogas*ter; At, *Arabidopsis thaliana*. **b** Interaction between LSM7 (wildtype) and LSM7-1t (truncated) with PCP, PCP-like and LSM5. pGADT7 and pGBKT7 bring AD and BD domains, respectively. After co-transformation, yeast cells were grown for six days on -WL at 28°C. The resulting colonies were then grown on both -WL and -WLH for four days at 28°C before taking photos. **c** A representative example of a T2 plant with ectopic LSM7 expression rescuing the embryo lethal phenotype of *lsm7-1* and showing abnormal leaf curvature grown at 23°C for 17 days.

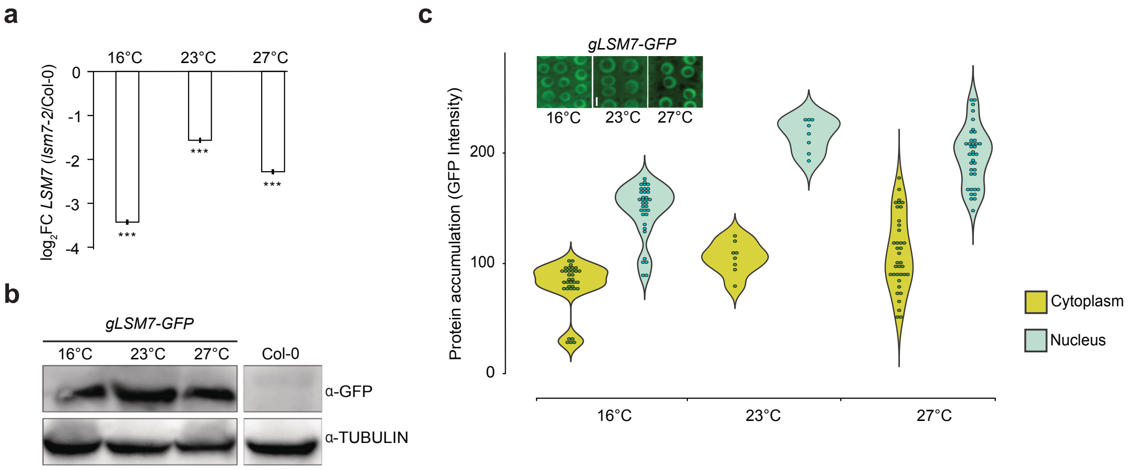


**Fig. S2. LSM7 distribution and stability are not affected by changes in ambient temperature.** **a** Relative *LSM7* expression ($log_2$FC) in different temperatures is shown as the ratio of *lsm7-2*/Col-0. The significance of FC values is indicated by three asterisks ($p<0.001$). **b** Accumulation of LSM7-GFP expressed under native promoter and terminator in plants grown under different ambient temperatures. Detection of LSM7-GFP by Western Blot using an anti-GFP antibody. Anti-TUBULIN is used as a loading control. **c** Subcellular localization of LSM7-GFP in plants grown at different temperatures. Each dot within the violin plots represents an independent measurement. Mean intensity indicates mean fluorescence signal intensity with 8-bit depth.

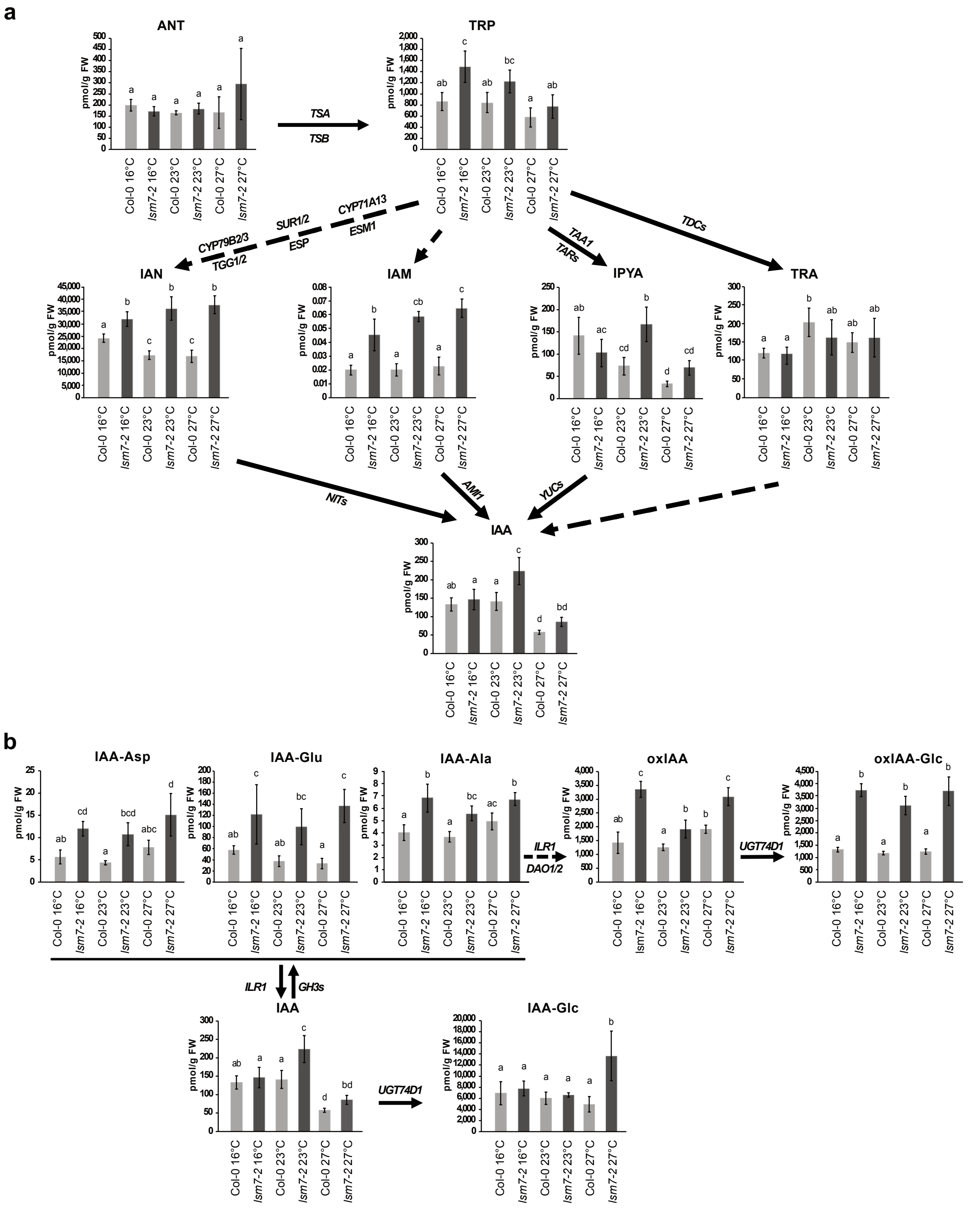


**Fig. S3. Auxin metabolites are modulated in the *lsm7-2* mutant.** Analysis of metabolites involved in (a) auxin biosynthesis and (b) auxin catabolism under different temperatures in the *lsm7*-2 mutant (dark grey) and Col-0 (light grey). Error bars represent the standard deviation (SD) of mean values from five biological replicates and letters of the statistical differences among the genotypes and conditions based on ANOVA and Tukey's HSD test. Quantification is presented as pmol/gram of fresh weight (pmol/g FW). Arrows indicate the main genes involved in the step. Solid lines indicate direct reactions, and dashed lines indicate indirect reactions.

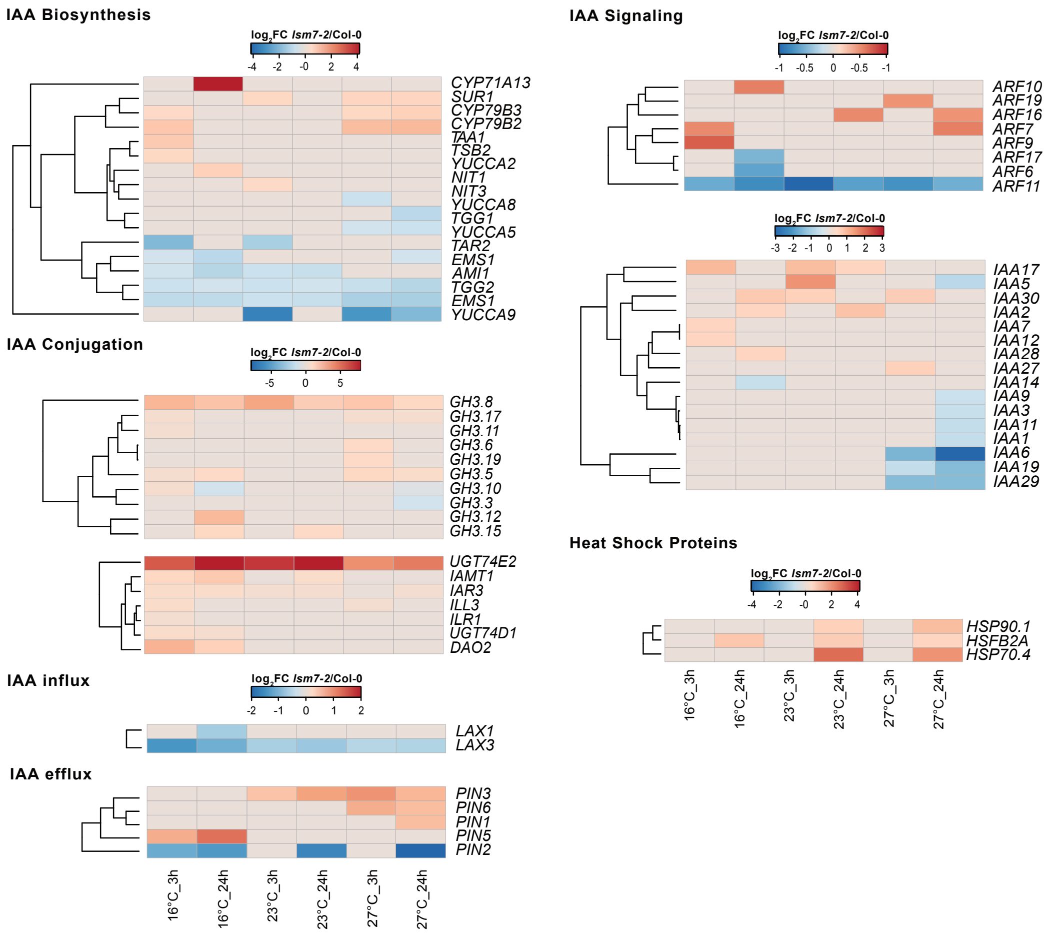


**Fig. S4. Auxin-related genes are heavily regulated *in lsm7-2* mutant.** Heatmaps of significant differentially expressed auxin-responsive genes' expression (log2FC *lsm7-2*/Col-0) in seedlings exposed to different ambient temperatures, organized according to their function. Shades of red indicate up-regulated genes, and shades of blue indicate downregulated genes in *lsm7-2.*

**Dataset S1 (separate file).** Differentially Expressed Genes (DE) under different treatments.

**Dataset S2 (separate file).** Differentially Alternatively Expressed (DAS) genes under different treatments.

**Dataset S3 (separate file).** Gene Ontologies (GOs) for Biological Processes (BP) of DE and DAS genes.

**Dataset S4 (separate file).** List of oligos and plasmids used in this study.
